# CRISPR Double Knockout Library Reveals Synthetic Lethal Gene Pairs in Triple-Negative Breast Cancer

**DOI:** 10.1101/2024.05.14.594157

**Authors:** Shuai Shao, Shangjia Li, Shan Tang, Kunjie Fan, Lang Li

## Abstract

Synthetic lethality, a genetic interaction involving two or more genes whose combined loss results in cell death, has emerged as a promising strategy for targeted cancer therapy. By exploiting synthetic lethal interactions, cancer cells can be selectively targeted and eradicated while preserving healthy cells, minimizing off-target effects, and reducing toxicity. The development of PARP inhibitors for ovarian and breast cancer patients with BRCA mutations exemplifies the potential of synthetic lethality-based therapy. Various experimental approaches, including CRISPR/Cas9 screens, have been employed to identify synthetic lethal gene pairs. Our lab has developed a CRISPR double knockout library, leveraging the XDeathDB database for candidate gene selection. This comprehensive platform offers insights into 12 cell death modes and 149 cell death hallmark genes. We aim to construct a cell-death double knock-out library using these genes and perform double knock-out screening on MDA-MB-231, a representative cell line for TNBC chemo poor responders. The identified synergistic lethal gene pairs may serve as potential drug targets for treating TNBC.

## Introduction

Synthetic lethality is a genetic interaction involving two or more genes, in which the combined loss of function or inhibition of these genes results in cell death, while the individual loss of function remains tolerable^1^. This concept has garnered considerable interest in recent years as a promising strategy for targeted cancer therapy. Exploiting synthetic lethal interactions between genes enables the selective targeting and eradication of cancer cells while preserving healthy cells, thus minimizing off-target effects, and reducing toxicity^2^.

A prime example of SL-based therapy is the development of PARP inhibitors for ovarian and breast cancer patients with BRCA mutations ^3, 4^. The synthetic lethal interaction between the BRCA and PARP genes led to the creation of drugs like niraparib, which selectively target cancer cells carrying BRCA mutations, sparing healthy cells. This breakthrough has revolutionized targeted cancer therapy, offering a more effective and less toxic treatment option for patients with BRCA-associated cancers ^5^. It also paves the way for further exploration of synthetic lethality in other genetic contexts to expand the range of targeted therapies against cancer.

Various experimental approaches, such as yeast two-hybrid systems ^6^, RNA interference (RNAi) screens ^7^, and genome-wide CRISPR-Cas9 screens^8, 9^, have been utilized to identify synthetic lethal gene pairs. Among these methods, the CRISPR/Cas9 system offers the greatest versatility for genetic manipulation. While RNAi functions at the post-transcriptional level, CRISPR/Cas9 is designed to generate functional knockouts at the gene level ^10–13^. Numerous research groups have compared RNAi- and CRISPR-based screening methods, demonstrating the higher efficiency of the CRISPR platform. These high-throughput techniques have expedited the discovery of many synthetic lethal interactions, with some successfully translating into clinical applications ^10–12^. In 2022, our lab developed a CRISPR double knockout library, improving screening efficacy by introducing two new backbones that offer options for screening cells with or without Cas9 expression modification ^14^.

Although CRISPR screening can effectively identify synergistic gene pairs, the capacity for combinatorial CRISPR screening is still constrained by library size and cell culture requirements. Even though the ability to screen gene combinations has grown, cell culture requires at least a few hundred times the size of the library, which places limitations on the cost of constructing double knockouts and the scale of cell capture. Consequently, the size of double knockout libraries that can be constructed is typically limited. Therefore, the selection of candidate genes remains critically important ^14^. In our lab’s previous research, we established the XDeathDB database^15^. This database can serve as a valuable resource for selecting candidate genes when constructing libraries. XDeathDB is acomprehensive bioinformatics platform designed to visualize and analyze the complex relationships between cell death processes and their crosstalk across 1461 cancer types. This platform consolidates data from various sources such as iRefIndex, STRING, BioGRID, Reactom, Pathway Commons, DisGeNET, DrugBank. By doing so, XDeathDB enables users to navigate extensive interdependent networks, identify hallmark genes, and uncover druggable targets within the cell death landscape, ultimately facilitating a deeper understanding of molecular mechanisms and promoting systems-oriented drug discovery.

XDeathDB offers a holistic perspective on 12 cell death modes, offering a detailed understanding of the complex interactions between these cell death modes.149 cell death hallmark (key) genes have been identified, providing valuable insights into the complex interactions among different cell death modes. The 149-cell death hallmark (key) genes have been experimentally validated to be significantly associated with cell death. Therefore, we aim to utilize these cell death hallmark genes as candidates for constructing a cell-death double knock-out library by overlapping our previous TKVO3 CRISPR screening and RNA-seq results. Subsequently, we will perform double knock-out screening on MDA-MB-231, the most representative cell line for TNBC poor responders that we have previously identified, to find synergistic lethal gene pairs for TNBC. These gene pairs can serve as potential drug targets for treating TNBC.

## Result

### Cell death CRISPR Double Knock Out construction

The candidate genes were selected by overlapping three sets of data, as shown in the Venn plot in Figure 2A. Through this process, 65 hallmark genes were identified for cell-death CDKO library construction, refining our selection among the 149 previously proven effective genes and focusing on those highly expressed in the MDA-MB-231 cell line while excluding single-gene lethal genes. The details of the 65 selected hallmark genes, including their gene names, functions, and relevant cell death mode, are presented in Table 2.

**Table 1.** CDKO sequencing statistic summary Table.

| **sample** | **Reads** | **mapped** | **Mapped**<br>**%** | **zero_**<br>**counts** | **zero_**<br>**counts%** | **Gini_**<br>**Index** | **above_**<br>**threshold** | **above_**<br>**threshold%** |
| --- | --- | --- | --- | --- | --- | --- | --- | --- |
| Library/Plasmid | 9272549 | 8159448 | 88.00 | 141 | 0.27 | 0.05 | 48921 | 92.48 |
| T0_1 | 12431419 | 11383720 | 91.57 | 892 | 1.69 | 0.06 | 47101 | 89.04 |
| T0_2 | 15647051 | 14281485 | 91.27 | 722 | 1.36 | 0.06 | 47669 | 90.11 |
| T0_3 | 15104601 | 13610181 | 90.11 | 1371 | 2.59 | 0.05 | 47547 | 89.88 |
| Tend_1 | 10004498 | 9114027 | 91.10 | 109 | 0.21 | 0.05 | 50658 | 95.76 |
| Tend_2 | 15386447 | 13994094 | 90.95 | 48 | 0.09 | 0.05 | 51875 | 98.06 |
| Tend_3 | 15837762 | 14264603 | 90.07 | 39 | 0.07 | 0.05 | 51866 | 98.05 |

**Table 2.** Hub genes with targeting inhibitor.

| Name | Pathway | Drug bank ID | Inhibitor Name | Previous Study in TNBC |
| --- | --- | --- | --- | --- |
| FTH1 | Autophagy | DB00852 | Pseudoephedrine | Prognostic Marker <sup>31</sup> |
| HDAC1 | Cell cycle | DB02546 | YF438 | anti-TNBC activity <sup>32</sup> |
| MAP3K3 | Efferocytosis | DB12010 | Fostamatinib | -- |
| DIABLO | Apoptosis | DB11752 | Bryostatin 1 | anti-TNBC activity <sup>33</sup> |
| VDAC2 | Ferroptosis | DB01375 | Aluminium monostearate | -- |
| CASP1 | Pyroptosis | DB00945 | Acetylsalicylic acid | anti-TNBC activity <sup>34</sup> |

**Table 3.**
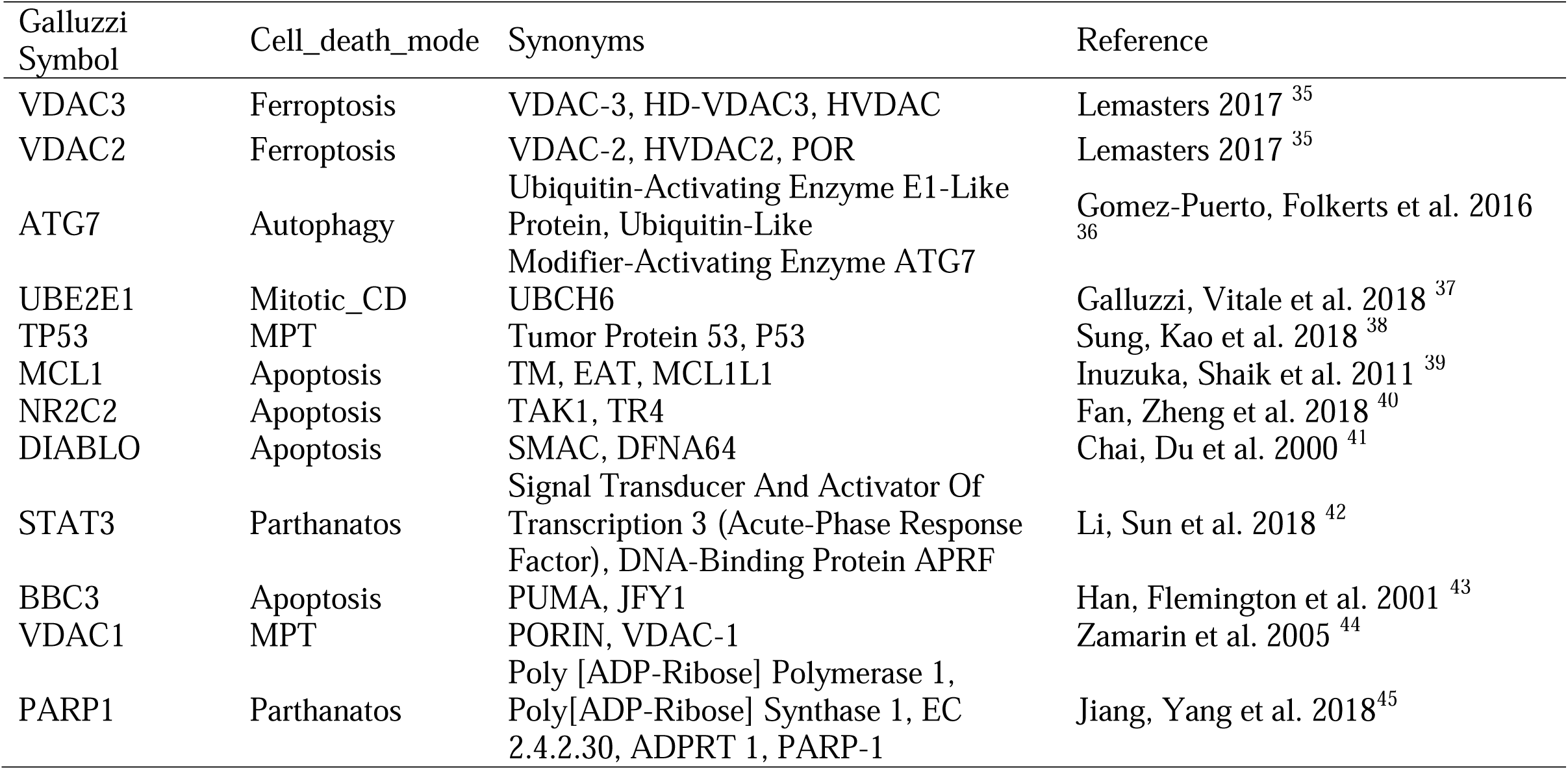
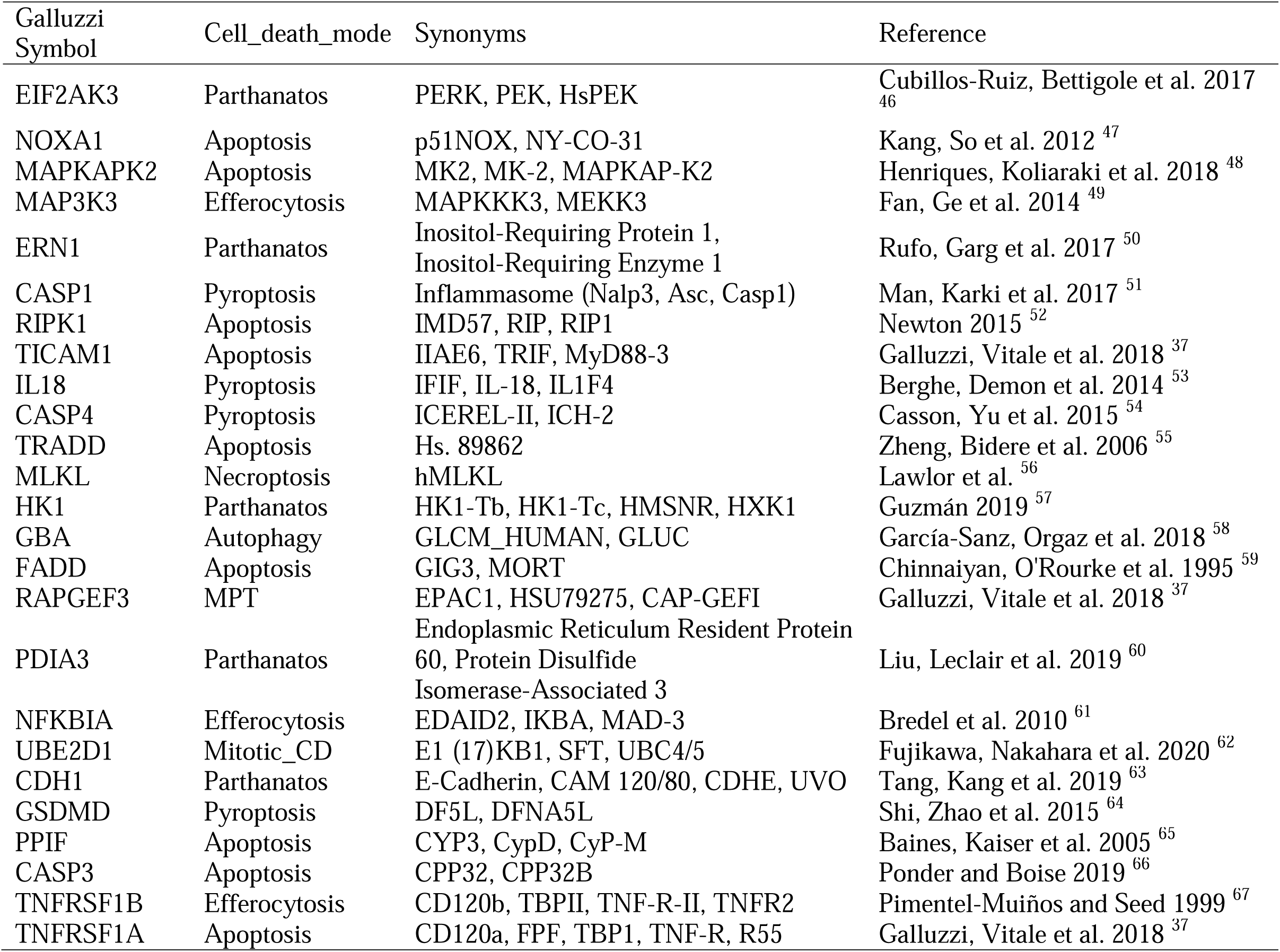
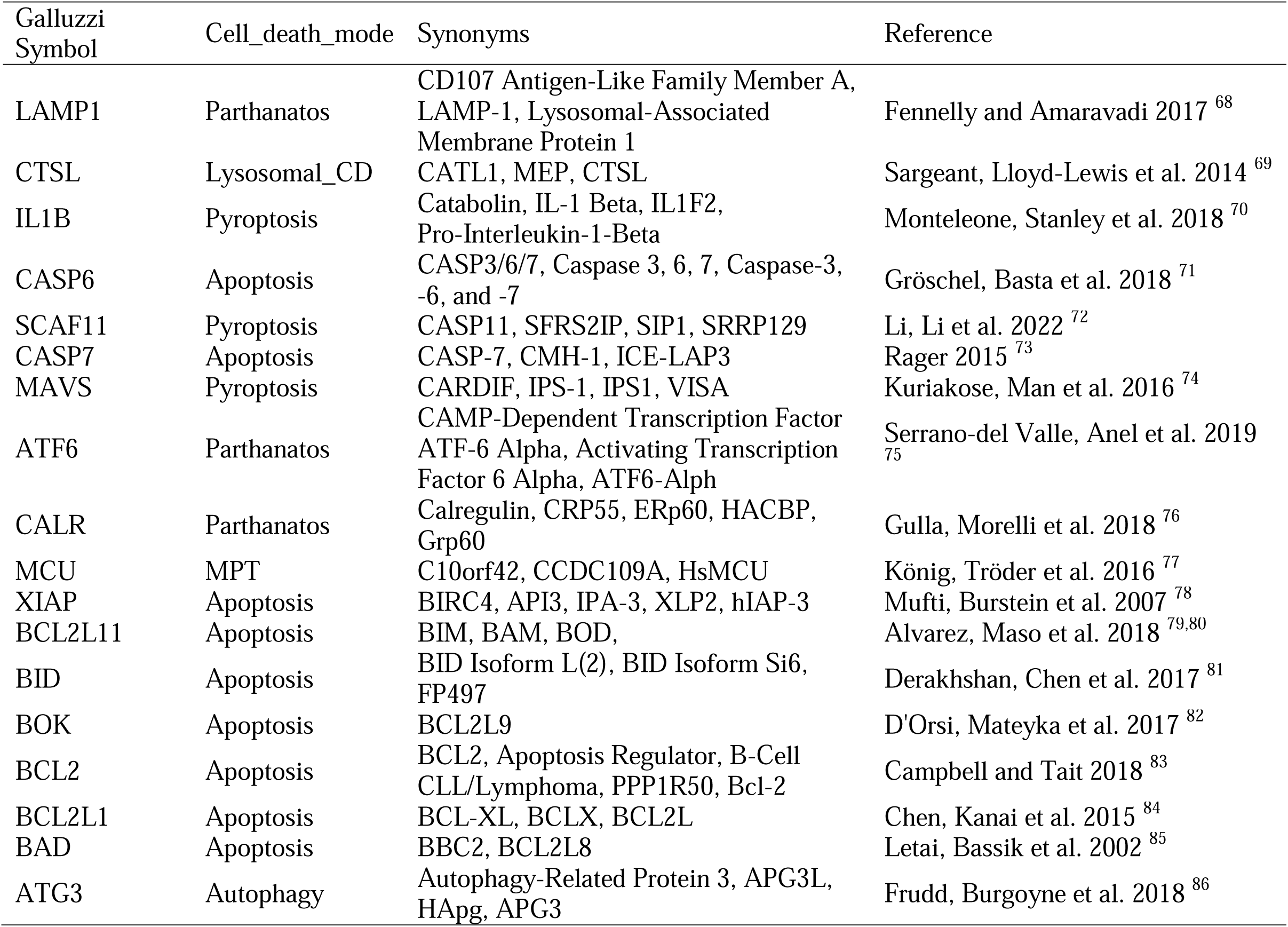
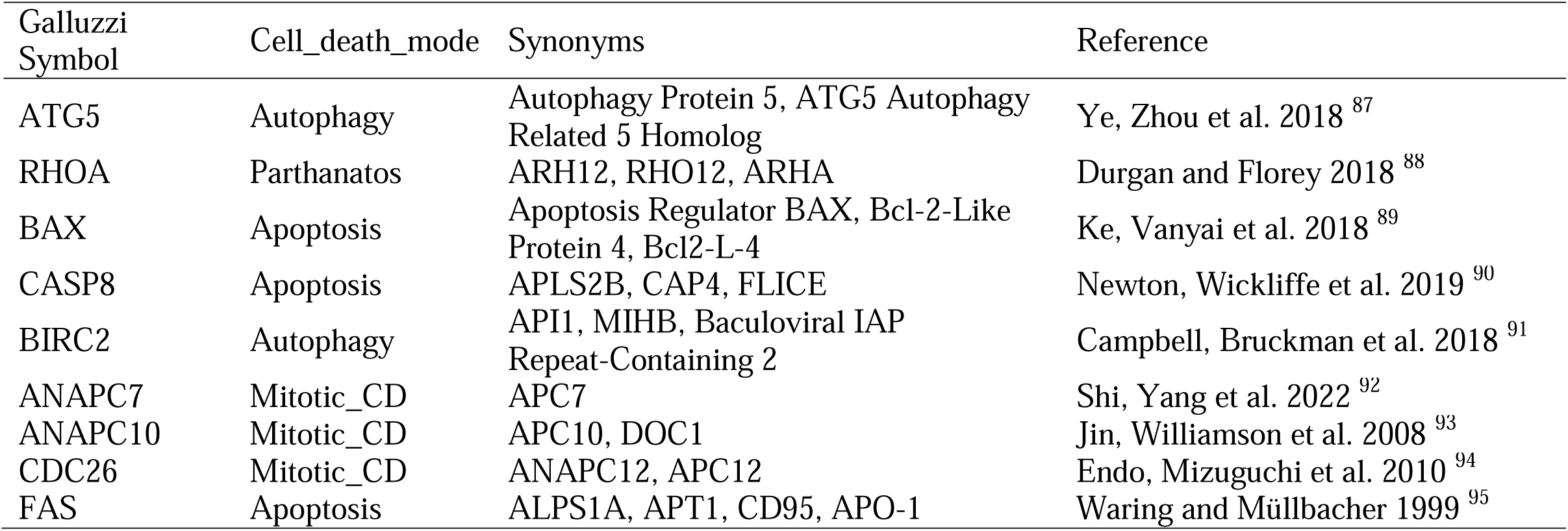
Candidate genes for developing Cell-death CDKO library.

### Pooled genome-wide CRISPR screening on Cas9-Expressing MDA-MB-231 Cell Line

After confirming the high quality of our cell death CDKO library, with only 141 missed gene-gene pairs and a low Gini index (Table 1), we conducted a genome-wide CRISPR/Cas9 pooled screen to identify synergistic lethal gene pairs in the MDA-MB-231 cell line (Figure 1), which is most representative cell line for TNBC poor responder. We transduced cells with the lentiviral pooled sgRNA library at a low MOI (0.3) and applied puromycin selection before collecting the baseline sample. After an additional 28 days of cell culture, we harvested triplicate samples. We performed deep sequencing and used multiple algorithms (see Methods) for data analysis. The sequencing results demonstrated high quality, with a mapping ratio of approximately 90% for all samples and around 10 million mapped reads per sample (Table 1, Figure 2B). The number of missed genes ranged from 39 to 1,000 sgRNA-sgRNA pairs out of a total of 52,900 sgRNAs (Table 1, Figure 2C). The Gini index for all samples was around 0.05 (Table 1, Figure 2D).

**Figure 1.**
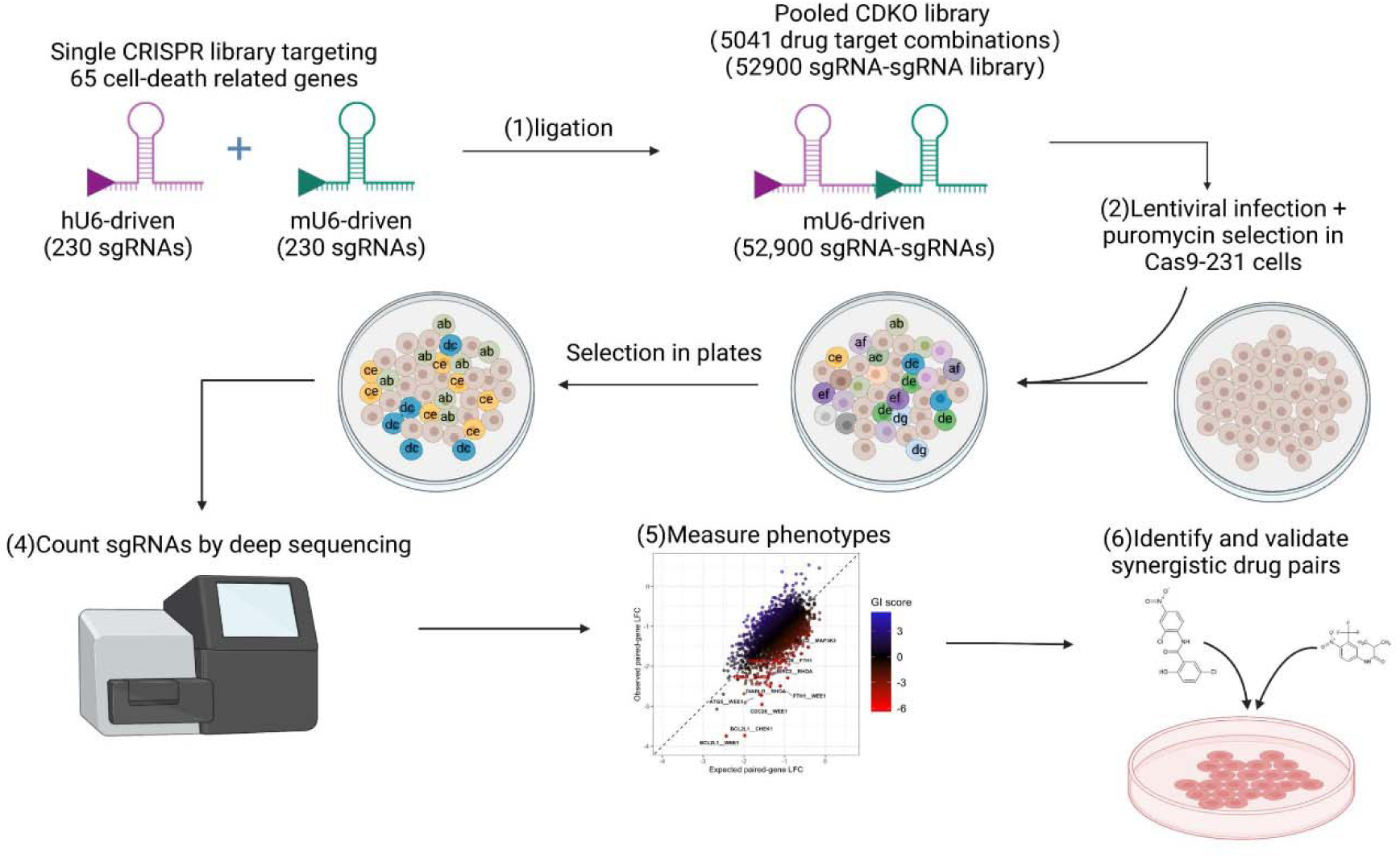
Schematic diagram for genome-wide CRISPR by the Cell-death CDKO library.

**Figure 2.**
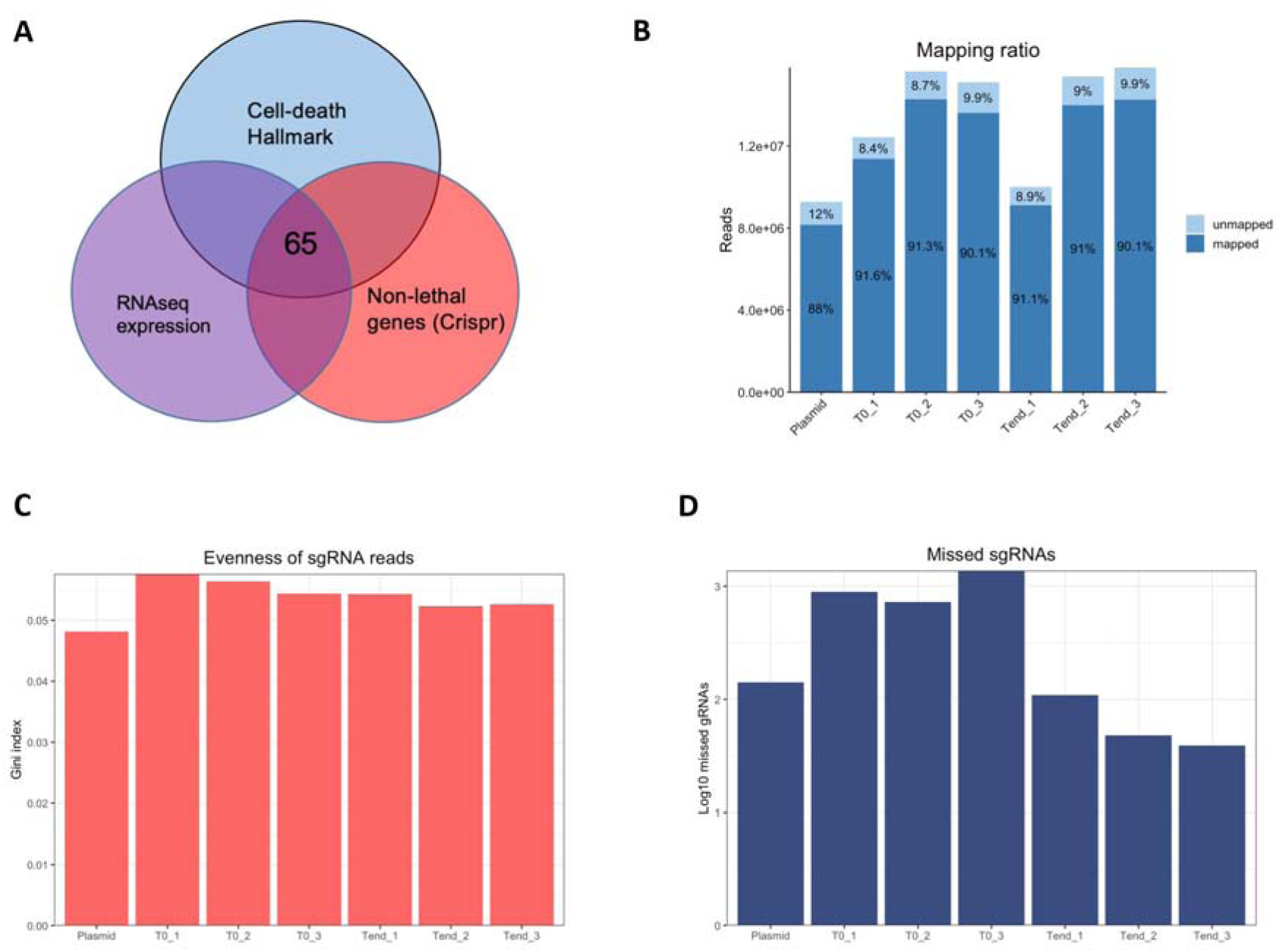
Cell-death Crispr CRISPR-Cas9 Double knock out screens in the MDA-MB-231 cell line. (A) Candidate gene selection for constructing cell-death CDKO library. (B) Read counts Mapping ratio of plasmid, days 0 (T0), and days 28 (Trend) samples. (C) The Gini index of sgRNAs on plasmid, days 0 (T0), and days 28 (Trend) samples. (D) The missed sgRNAs were tested on plasmid, days 0 (T0), and days 28 (Trend) samples.

### Essential gene identification

From the PCA plot (Figure 3A) and pairwise sample correlation plot (Figure 3B), it is evident that samples within the same treatment group demonstrate a distinct pattern and higher correlation values. This observation indicates consistent results among triplicate samples. Overall, the data analysis supports the successful implementation of our CRISPR-based screen in the MDA-MB-231 cell line. We identified potential hit gene pairs by comparing the observed LFCs for gene-gene pairs with the expected LFCs derived from individual gene LFCs (Figure 3C-3D). Moreover, we employed advanced calculation methods based on seven different approaches (see Methods) for SL gene pair selection. The top 10% pairs (248 pairs) from each scoring method were chosen, and their overlaps are depicted in both the Venn plot and histogram (Figure 4A and 4B). 242 gene pairs (∼10%) exhibited synthetic lethality in more than three methods, and these pairs are more likely to have an SL effect compared to other pairs. We constructed a gene-gene network based on these 242 gene pairs (Figure 4C). Hub genes interacting with at least ten other genes are displayed in the histogram (Figure 4D).

**Figure 3.**
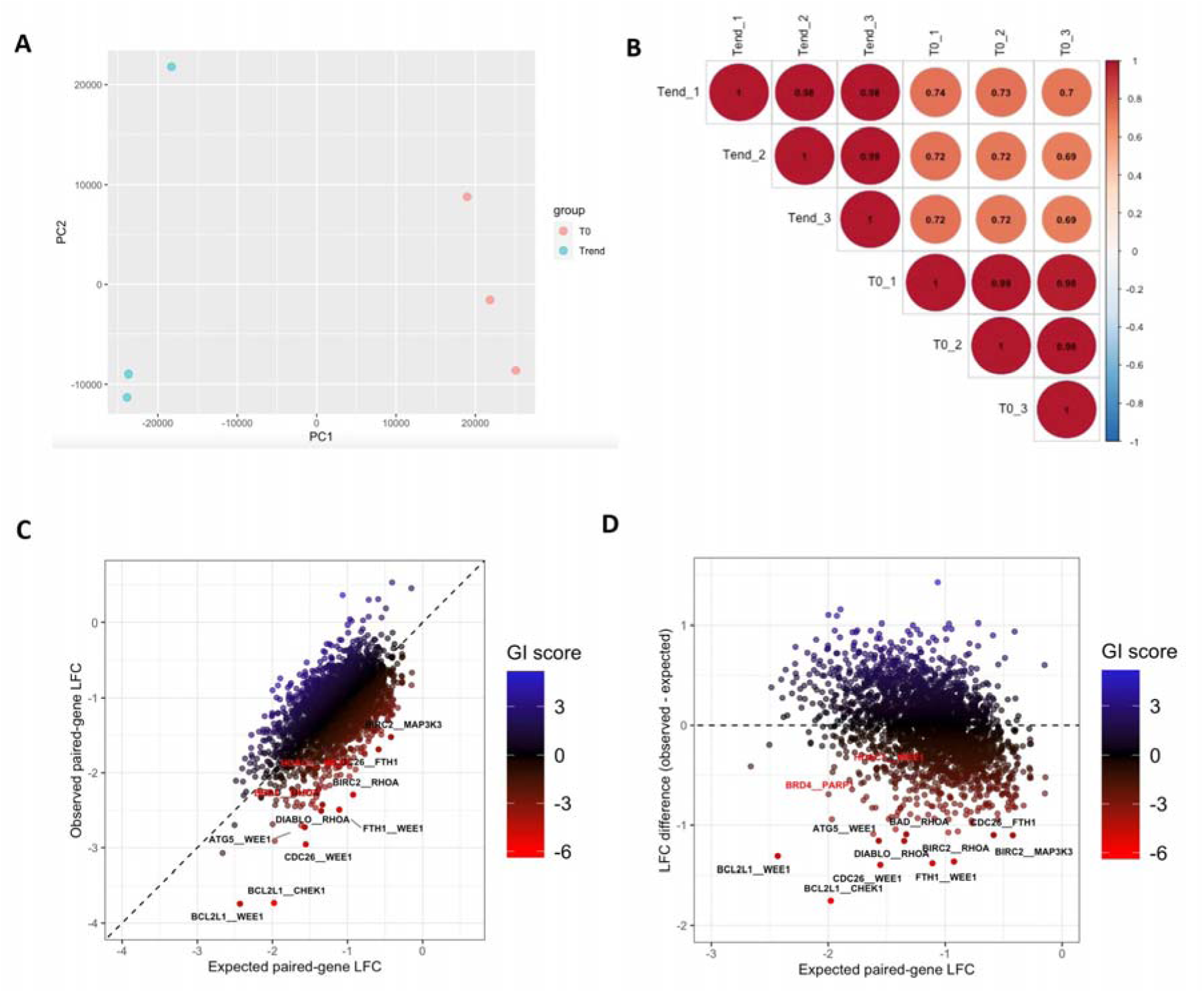
CDKO data exploration and synergistic lethal paired-gene LFC visualization. (A) PCA plot of baseline(T0) and Day28(Trend) samples. (B) Correlation plot between baseline(T0) and Day28(Trend) group, each group containing triplicate samples. LFC deviation between observed and expected gene-gene pairs (C) Observed LFC vs Expected LFC (D) Difference between observed LFC and Expected LFC vs Expected LFC.

**Figure 4.**
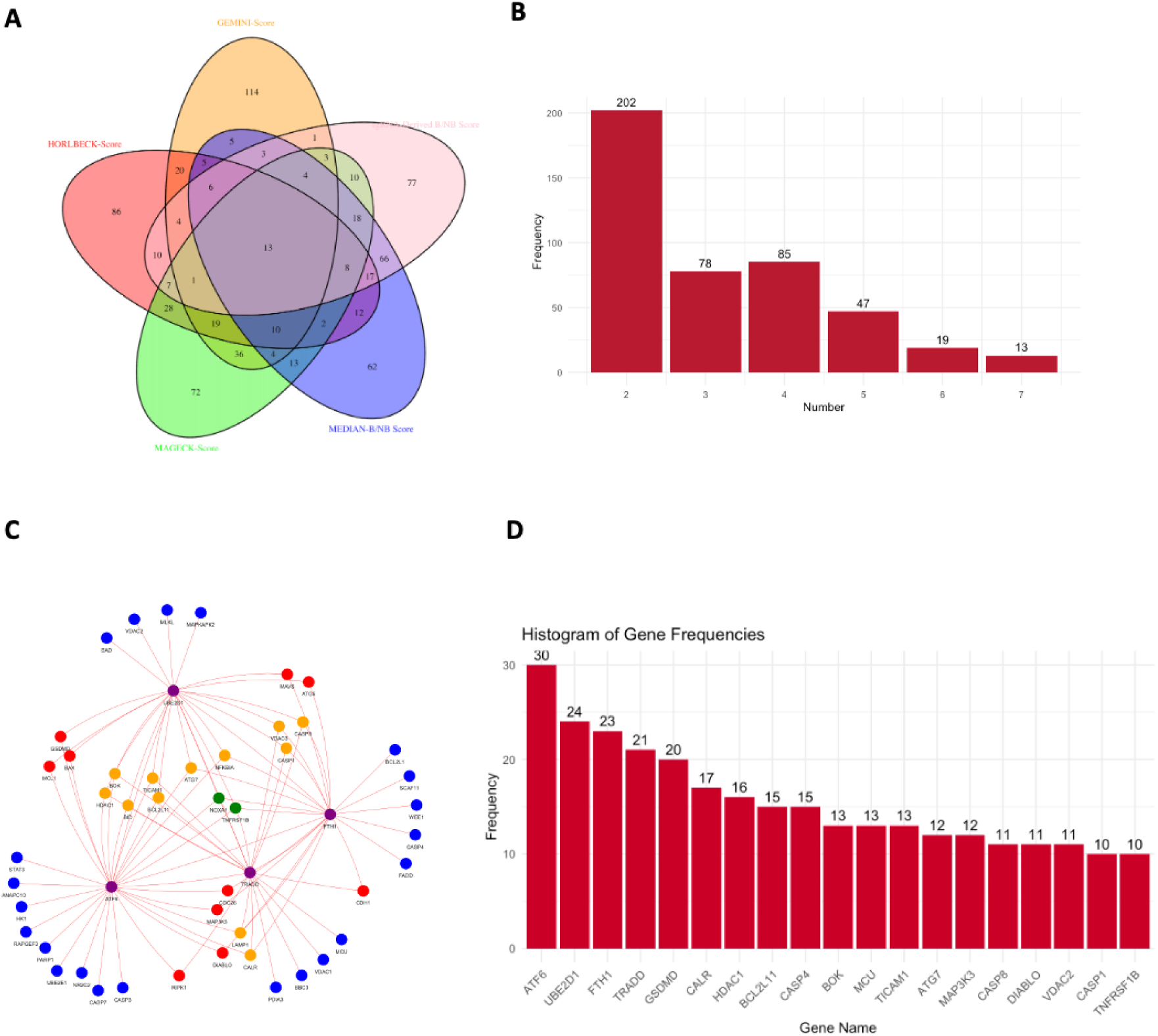
Synergistic lethal gene pair identification. (A) Venn Plot of 7 SL pairs scoring methods. (B) Histogram displaying the frequency distribution of SL gene pairs overlapping results among 7 different methods. (C) Gene network by 242 SL plots (D) Histogram displaying the frequency distribution of top hub genes in 242 SL gene pairs.

## Discussion

In this study, we successfully applied a genome-wide CRISPR/Cas9 pooled screen using our cell death CDKO library to identify synthetic lethal gene pairs in the MDA-MB-231 cell line, a representative model for TNBC poor responders. Our results reveal 135 synthetic lethal pairs, providing potential novel therapeutic targets for the treatment of TNBC.

The high quality of construction of our cell death CRISPR-based screen library was supported by the low number of missed gene-gene pairs, the high mapping ratio, and the low Gini index for all samples. The consistency in the results observed in triplicate samples further validates our experimental approach. The use of advanced calculation methods, such as sgRNA median score and sgRNA derived score, in combination with stringent selection criteria, enabled us to pinpoint the top 10% gene pairs with the highest likelihood of synthetic lethality.

In our study, we introduced five synthetic lethal gene-gene pairs as positive controls. Based on the results from our screening data analysis, four out of the five gene pairs were ranked within the top 10% according to their LFC ranking scores: BRD4 PARP1 with a rank of 9 and an SL score of 8.56E-10, HDAC1 WEE1 with a rank of 124 and an SL score of 3.29E-05, BRD4 CHEK1 with a rank of 172 and an SL score of 0.00016564, and HDAC2 WEE1 with a rank of 200 and an SL score of 0.00027904. The presence of most positive gene-gene pairs in our results demonstrates that our experiment is successful and the outcome is reliable. HDAC1 inhibition has been shown to induce histone tail acetylation and an open chromatin structure, which promotes the expression of cell differentiation or death genes. In combination with WEE1 inhibition, the impaired cell cycle checkpoint kinase activity and premature mitotic entry lead to an increase in DNA damage^16^ and apoptosis, making this combination a promising target for the treatment of AML^17^. Similarly, Inhibition of BRD4 has been shown to induce HRD and sensitize cells to PARP inhibitors. This combination has the potential to reverse intrinsic resistance in RAS mutant tumors and other mechanisms of PARPi resistance, warranting clinical assessment in both PARPi-sensitive and -resistant cancers^18^.

Our findings add to the ongoing efforts to discover novel synthetic lethal interactions that can be harnessed for targeted cancer therapy. The 242 synthetic lethal pairs identified in this study offer a valuable resource for future research in TNBC and other cancer types. Among the 242 gene pairs we identified, some have already been reported to demonstrate synthetic lethality effects. CDH1-HDAC1 and HDAC1-PARP1 gene pairs have been reported as synergistically lethal gene pairs and can be considered potential drug targets. Decourtye-Espiard et al. investigated the potential of histone deacetylase (HDAC) inhibitors as a treatment for cancers with CDH1 mutations, testing the effects of several HDAC inhibitors on gastric and breast preclinical models with and without CDH1 mutations^19^. They found that CDH1-null cells were more sensitive to pan-HDAC inhibitors such as entinostat, pracinostat, mocetinostat, and vorinostat compared to wild-type cells. This supports the notion that CDH1-HDAC1 has a strong synergistic lethal effect on cancer. The combination of hypomethylating agents (HMAs), inhibitors of poly(ADP-ribose) polymerases (PARPis), and histone deacetylases (HDACis) was hypothesized to be synergistically cytotoxic to leukemia and lymphoma cells. Valdez BC et al. demonstrated that exposing AML and lymphoma cell lines to a combination of PARPi niraparib, HMA decitabine, and HDACi romidepsin or panobinostat led to a synergistic inhibition of cell proliferation by up to 70%. This combination activated the ATM pathway, increased the production of reactive oxygen species, decreased mitochondrial membrane potential, and induced apoptosis^19^. This provides robust evidence that our CRISPR double knockout experiments are indeed effective in identifying synthetic lethal gene pairs.

In the gene-gene network focusing on 242 SL interactions, we specifically select hub genes of interest, which have the highest number of pairs (Figure 4 D). The shared network employs a color-coding scheme to visualize connections between these targeted genes (purple) and their interacting partners: blue for 1 connection, red for 2 connections, yellow for 3 connections, and green for 3 or more connections with the targeted genes. Among the 19 hub genes, 6 of them were found have related drug inhibitors, most of them have been already identified in previous research for TNBC treatment (Table2). Although individual drugs have been studied, there have been no research reports on drug combination studies. This provides us with ample opportunity to discover the most novel and effective drug combinations for treating triple-negative breast cancer (TNBC).

Although our study successfully identified synthetic lethal gene pairs in the MDA-MB-231 cell line, further validation of these interactions in additional TNBC cell lines and in vivo models is necessary to confirm their relevance and potential as therapeutic targets. To achieve this, we need to conduct more validation experiments, such as designing a CRISPR screening library that targets only the 242 known gene pairs and performing two-drug combination assays. Additionally, we require detailed functional characterization of the identified gene pairs to understand the underlying molecular mechanisms driving synthetic lethality and determine the optimal drug combinations and dosages for clinical translation.

In conclusion, our CRISPR-based screening approach has successfully identified synthetic lethal gene pairs in a representative TNBC cell line, offering new potential drug targets for the treatment of this aggressive and challenging cancer subtype. Future studies should focus on validating these interactions in additional models and characterizing their functional roles to pave the way for the development of innovative, targeted therapies for TNBC patients.

## Method

### Synthetic lethal gene candidates’ selection

To select synthetic lethal gene candidates for constructing a CDKO CRISPR library, we refined our selection among 149 hallmark genes that were previously proven effective. Our screening criteria involved two main steps: (1) selecting genes that were highly expressed in the MDA-MB-231 cell line by referring to RNA-sequencing data from previous experiments and identifying genes with read counts greater than 50 as highly expressed genes (2) excluding lethal genes causing single-gene lethality in the MDA-MB-231 cell line. We leveraged data from our previous TKOV3 CRIPSR Cas9 screening to identify 1400 non-lethal genes in the MDA-MB-231 cell line. By overlapping these three sets of data, we ultimately screened a total of 65 hallmark genes as synthetic lethal gene candidates.

### Library construction

The cell-death CRISPR Double knock out (CDKO) screening library was developed following the protocol our lab published in 2022 ^20^. To ensure the screening quality, we required three sgRNAs per gene. In our library, three corresponding sgRNAs were selected for each of the 65 genes, plus six positive control genes (71 genes in total), by referencing widely-used CRISPR libraries, such as TKOv3 ^21^, hGECKOv2 ^22^, and KinomeKO/Brunello ^23^, We also considered the VBC (Vienna Bioactivity CRISPR) scores ^24^ for each sgRNA. Additionally, we included 17 safe sgRNAs (8% of the total) that target non-functional regions of the genome as negative controls in the pooled library.

The cell-death CDKO library, comprising 52,900 sgRNAs targeting 5,041 gene-gene pairs, was developed and amplified 1,000-fold using the electroporation method. For lentivirus production, 5 million 293T cells were seeded in 15 cm plates and prepared for transfection. The packaging vectors psPAX2, pMD2.G (Addgene), the cell-death CDKO library plasmid, and Lipofectamine (Thermo Fisher Scientific) were combined in OptiMEM (Thermo Fisher Scientific, Waltham, MA, USA). After 48 hours of incubation, the lentivirus-containing medium was harvested and stored at −80°C.

### Cell Culture

The MDA-MB-231 human triple-negative breast cancer cell line and the 293T Homo sapiens embryonic kidney cell line, procured from the American Type Culture Collection (Manassas, VA, USA), were utilized in this study. Cas9-Expressing on MDA-MB-231 Cell Line were developed in Dr. Parvin’s lab. Each cell line was cultured in Ham’s F-12K (Kaighn’s) medium, enriched with 10% fetal bovine serum (VWR, Radnor, PA, USA), 1% GlutaMAX, 1% sodium pyruvate, and a penicillin-streptomycin mix (Gibco, Waltham, MA, USA). The cells were incubated at 37 °C in an environment containing 5% CO2. To ensure the integrity of the cell lines, they were subjected to STR profiling for authentication, and mycoplasma contamination tests were performed every three months.

### Pooled sgRNA screens

MDA-MB-231 cells were infected with the cell death CDKO lentiviral library at a low MOI of 0.3. Following 72 hours of 5μg/mL puromycin treatment, the surviving cells were designated as baseline samples (T0), and 3×10^7 cells were gathered and preserved at −80°C. The remaining cells were split into three separate groups. After a 28-day cultivation period, 3×10^7 cells were collected from each group (Trend). Genomic DNA was isolated using the QIAamp Blood Maxi Kit (Qiagen, Hilden, Germany). A series of two polymerase chain reactions (PCRs) were conducted to enrich sgRNA-targeted genomic regions and amplify the sgRNA. The derived libraries were sequenced on a NextSeq 500 system (Illumina, San Diego, CA, USA), generating nearly 10 million reads per sample and attaining 200x coverage of the cell death CDKO library.

### CDKO Crispr Data analysis

To analyze the data from the cell-death CDKO CRISPR screen, we first used FastQC to obtain an overview of basic quality control metrics for the raw next-generation sequencing data. Next, the dual-crispr tool was employed to map and count the gRNA-gRNA constructs. We then calculated the log fold change (LFC) for the abundance of each gene pair using MAGeCK RRA, pooling dual-gRNA constructs targeting the same gene pair together. ^25^ Then, we calculated the genetic interaction (GI) scores and identified potential hit gene pairs by comparing observed LFCs for gene-gene pairs to expected LFCs based on individual gene LFCs.^20^ To predict synthetic lethal pairs, we employed two SL scoring methods utilizing log2 fold change (LFC): sgRNA derived score ^26^and direct median score^27^. Following counts normalization, SL scores were calculated by taking the difference of expected to the observed sgRNA LFC, where expected sgRNA LFC is the sum of LFCs of sgRNAs paired with controls, negative scores indicating synthetic lethal activity and positive scores indicating buffering. Both scores were both normalized to the paired sgRNAs targeting controls by reducing the LFCs by median of control pair LFC.

### Synergistic lethal gene pair score calculations by using multiple methods

GEMINI: The R package GEMINI (v. 1.12.0) ^28^was employed to compute scores, using default settings that included normalization and a pseudo count of 32. The model was built with control targeting sgRNAs, followed by the package’s fitting process. The highest scores, indicative of the strongest interactions, were utilized to predict synthetic lethal pairs.

Horlbeck-Score: This scoring technique is adapted from Horlbeck et al. 2018^29^. Gene interaction scores were determined across sgRNA pairs at each orientation through a quadratic fit of sgRNA log fold change (LFC) counts. The synthetic lethal score is the difference between the fit and the observed LFC, with the most lethal pairs having the most negative scores. Lethality is represented by negative scores.

MAGeCK-Score: The MAGeCK script was used to calculate LFC values for sgRNA gene pairs with default parameters ^30^. The expected LFC was then computed by summing the LFC of each targeted gene pair and control. The synthetic lethal scores were calculated by comparing the expected and observed sgRNA pair LFC at the gene level using the median. Lethality is indicated by negative scores.

Median-B/NB Score: In this method, which is based on LFC synthetic lethal score calculations, sgRNA counts are first normalized across various time points. The LFC is then computed using the median of experimental replicates at the initial and final time points of the CRISPR study. Like the MAGeCK-Score, the expected LFC is calculated and used to determine the synthetic lethal scores. The standard error is calculated for each gene pair and employed to compute z-scores. Lethality is represented by negative scores.

sgRNA Derived Score B/NB: Following the LFC synthetic lethal score calculation methods, this score is determined based on each replicate. The LFC is computed for each experimental replicate across the initial and final time points. Next, the replicate SL score is calculated by taking the median of the difference between the expected and observed sgRNA pair LFC at the gene level. Similar to the Median-B/NB method, standard error is used to derive z-scores. The final scores for the study are represented by the median of replicate z-level SL scores. Lethality is indicated by negative scores.

